# Dissociating the neural correlates of subjective visibility from those of decision confidence

**DOI:** 10.1101/2021.06.01.446101

**Authors:** Matan Mazor, Nadine Dijkstra, Stephen M. Fleming

## Abstract

A key goal of consciousness science is identifying neural signatures of being aware vs. unaware of simple stimuli. This is often investigated in the context of near-threshold detection, with reports of stimulus awareness being linked to heightened activation in a frontoparietal network. However, due to reports of stimulus presence typically being associated with higher confidence than reports of stimulus absence, these results could be explained by frontoparietal regions encoding stimulus visibility, decision confidence or both. In an exploratory analysis, we leverage fMRI data from 35 human participants (20 females) to disentangle these possibilities. We first show that, whereas stimulus identity was best decoded from the visual cortex, stimulus visibility (presence vs. absence) was best decoded from prefrontal regions. To control for effects of confidence, we then selectively sampled trials prior to decoding to equalize confidence distributions between absence and presence responses. This analysis revealed striking differences in the neural correlates of subjective visibility in prefrontal cortex regions of interest, depending on whether or not differences in confidence were controlled for. We interpret our findings as highlighting the importance of controlling for metacognitive aspects of the decision process in the search for neural correlates of visual awareness.

**Significance statement:** While much has been learned over the past two decades about the neural basis of visual awareness, the role of the prefrontal cortex remains a topic of debate. By applying decoding analyses to functional brain imaging data, we show that prefrontal representations of subjective visibility are contaminated by neural correlates of decision confidence. We propose a new analysis method to control for these metacognitive aspects of awareness reports, and use it to reveal confidence-independent correlates of perceptual judgments in a subset of prefrontal areas.

## Introduction

In neuroimaging studies of visual perception, frontal and parietal cortices typically show stronger activation when participants report being aware rather than unaware of a visual stimulus (Sahraie et al., 1997; Dehaene et al., 2001; Fisch et al., 2009; Koivisto & Revonsuo, 2010). This finding is a cornerstone of several influential theories of awareness (e.g., *Global Neuronal Workspace*: Dehaene, Sergent & Changeux, 2003; Dehaene., Changeux, & Naccache, 2011; *Higher Order Thought*: Lau & Rosenthal, 2011; Brown, Lau, & LeDoux, 2019), and is central to recent debates about the specific role of these regions in the generation of subjective experience (Boly et al., 2017; Odegaard, Knight & Lau, 2017; Michel & Morales, 2020; Raccah, Block & Fox, 2021).

However, reports of awareness and unawareness of a visual stimulus differ not only in terms of whether a stimulus was visible or not, but also in other cognitive factors (Bayne & Hohwy, 2013). Specifically, when asked to rate their subjective confidence in near-threshold detection, participants’ confidence in decisions about stimulus presence is reliably higher than in decisions about stimulus absence (Mazor, Friston & Fleming, 2020; Mazor, Moran & Fleming, 2021). This confidence asymmetry between judgments of presence and absence makes interpreting frontoparietal activations in reports of visual awareness difficult: they may reflect stimulus visibility, subjective confidence in the percept (which is higher when a stimulus is detected), or both.

Consistent with the idea that frontoparietal activations found to correlate with awareness might reflect confidence, the same regions associated with awareness reports are also found to be implicated in reports of subjective confidence. For example, a coordinate-based meta-analysis revealed that dorsolateral prefrontal cortex, lateral parietal cortex, and posterior medial frontal cortex show a reliable parametric modulation of confidence (Vacarro & Fleming, 2018) -all regions that have been associated with subjective visibility in previous studies (Sahraie et al., 1997; Dehaene et al., 2001; Lau & Passingham, 2008; Fisch et al., 2009; Koivisto & Revonsuo, 2010). Importantly, these regions encode subjective confidence not only in perceptual decisions, but also in memory-based (Morales, Lau & Fleming, 2018) and value-based (De-Martino et al., 2013) decisions, suggesting that their link to subjective confidence is not solely in virtue of their role in tracking subjective visibility.

Here, we set out to systematically dissociate the neural correlates of visibility and confidence, and ask to what extent neural representations within a frontoparietal network track one or both of these variables. To address this question, we performed a series of exploratory analyses on neuroimaging data collected during performance-matched visual detection and discrimination tasks with subjective confidence ratings (originally reported in Mazor et al., 2020). We first asked where in the brain can we decode the presence or absence of a visual target stimulus (a sinusoidal grating) from multivariate spatial activity patterns during the detection task. By comparing these results against similar decoding of stimulus identity (grating orientation) in a performance-matched discrimination task, we could control for non-specific neural contributions to perceptual decision-making and report. Critically, by leveraging trial-wise confidence ratings we were able to equate differences in subjective confidence between conditions, allowing us to isolate neural representations associated with stimulus visibility. To anticipate our results, we find that a number of prefrontal representations of stimulus visibility are confounded with representations of confidence, but that a confidence-independent representation of perceptual content is present in posterior medial frontal cortex (pMFC). Our approach provides a novel method for controlling for such confidence effects in future studies of visual awareness.

## Methods

This is an exploratory analysis of neuroimaging data, originally reported in Mazor et al. (2020). For a more elaborate description of the experimental design and behavioural findings, see Mazor et al. (2020).

### Participants

46 participants took part in the study (ages 18–36, mean = 24 ± 4). We applied the same subject- and block-wise exclusion criteria as in the original study. Specifically, participants were excluded for having low response accuracy, pronounced response bias, or insufficient variability in their confidence ratings. 35 participants met our pre-specified inclusion criteria (ages 18–36, mean = 24 ± 4; 20 females). We pre-registered a sample size of 35 to maximize statistical power given resource limitations. This allowed us to detect a medium effect in a paired-samples t-test (cohen’s d = 0.49) with a power of 80%. All analyses are based on the included blocks from these 35 participants.

Pre-registration was time-locked by initializing the pseudorandom number generator with a hash of our pre-registered protocol folder (link: github.com/matanmazor/detectionVsDiscrimination_fMRI/tree/master/protocol folder) prior to determining the order and timing of experimental events (Mazor, Mazor & Mukamel, 2019).

Importantly, this pre-registration was motivated by a different set of hypotheses (tested in Mazor, Friston & Fleming, 2020). The results we present here are derived from a data-driven, exploratory set of analyses.

## Experimental Design and Statistical Analysis

### Design and procedure

Trials started with a fixation cross (500 milliseconds), followed by a presentation of a stimulus for 33 milliseconds. In discrimination trials, the stimulus was a circle of diameter 3° containing randomly generated white noise, merged with a sinusoidal grating (2 cycles per degree; oriented 45° or −45°). In half of the detection trials, stimuli did not contain a sinusoidal grating and consisted of random noise only. After stimulus offset, participants used their right-hand index and middle fingers to make a perceptual decision about the orientation of the grating (discrimination blocks), or about the presence or absence of a grating (detection blocks; see Fig. 1, top panel). Response mapping was counterbalanced between blocks which means that significant decoding of decisions cannot reflect motor representations.

**Figure 1:**
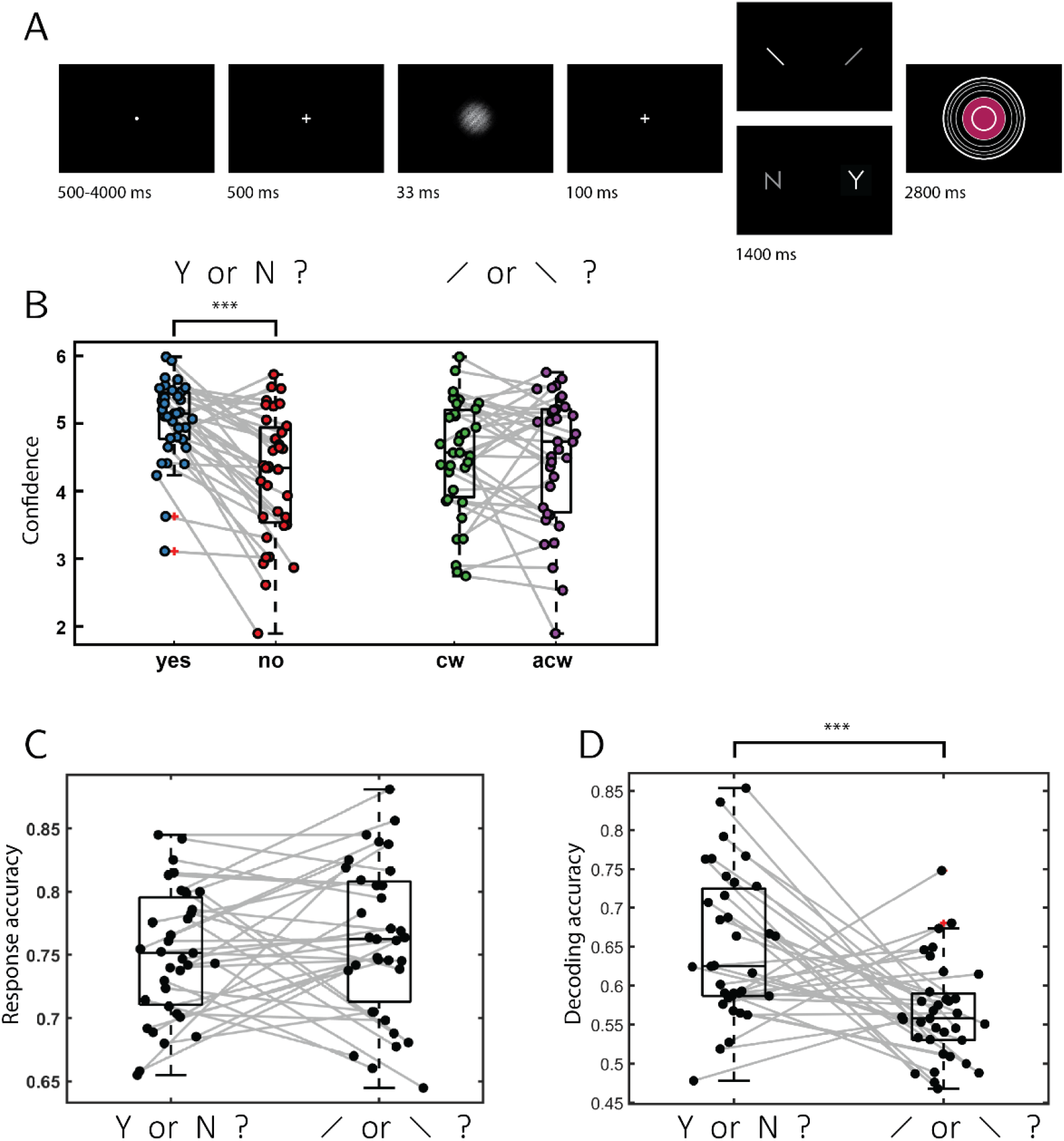
Experimental design and behavioural results. A: In discrimination trials, participants made discrimination judgments about clockwise and anticlockwise tilted noisy gratings, and then rated their subjective confidence by controlling the size of a colored circle. In detection judgments, decisions were made about the presence (Y) or absence (N) of a grating in noise. B: mean confidence as a function of response for the 35 participants. Confidence in detection ‘yes’ responses was significantly higher than in ‘no’ responses. No significant difference was observed between confidence in discrimination responses (cw: clockwise, acw: anticlockwise). C: Response accuracy was not different between the two tasks. D: Decoding accuracy for a classifier trained to classify response (yes or no in detection, clockwise or anticlockwise in discrimination) based on confidence ratings alone. Decoding accuracy was significantly higher for detection than for discrimination. ***: p<0.001.

Immediately after making a decision, participants rated their confidence on a 6-point scale by using two keys to increase or decrease their reported confidence level with their left-hand thumb. Confidence levels were indicated by the size and color of a circle presented at the center of the screen. The initial size and color of the circle was determined randomly at the beginning of the confidence rating phase. The mapping between color and size to confidence was counterbalanced between participants: for half of the participants high confidence was mapped to small, red circles, and for the other half high confidence was mapped to large, blue circles. The perceptual decision and the confidence rating phases were restricted to 1500 and 2500 milliseconds, respectively. No feedback was delivered to subjects about their performance. Trials were separated by a temporally jittered rest period of 500-4000 milliseconds.

Prior to the scanning day, participants underwent a behavioral session in which task difficulty was adjusted independently for the detection and discrimination tasks, targeting around 70% accuracy. We achieved this by adaptively adjusting the stimulus signal-to-noise ratio (SNR) every 10 trials (increasing the SNR if accuracy fell below 60%, and decreasing it if accuracy exceeded 80%). Task difficulty was further calibrated within the scanner environment at the beginning of the scanning session, during the acquisition of anatomical (MP-RAGE and fieldmap) images, using a similar procedure. Upon completion of the calibration phase, participants performed 5 experimental runs comprising one discrimination and one detection block, each of 40 trials, presented in random order. A bonus was awarded for accurate responses and confidence ratings according to the following formula: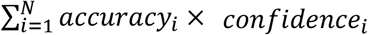, where *accuracy* equals 1 for correct responses and -1 for incorrect responses, and *confidence* is the reported confidence level on a scale of 1-6.

### Scanning parameters

Scanning took place at the Wellcome Centre for Human Neuroimaging, London, using a 3 Tesla Siemens Prisma MRI scanner with a 64-channel head coil. We acquired structural images using an MPRAGE sequence (1×1×1 mm voxels, 176 slices, in plane FoV = 256×256 mm2), followed by a double-echo FLASH (gradient echo) sequence with TE1 = 10 ms and TE2 = 12.46 ms (64 slices, slice thickness = 2 mm, gap = 1 mm, in plane FoV = 192 × 192 mm2, resolution = 3 × 3 mm2) that was later used for field inhomogeneity correction. Functional scans were acquired using a 2D EPI sequence, optimized for regions near the orbito-frontal cortex (3×3×3 mm voxels, TR = 3.36 s, TE = 30 ms, 48 slices tilted by −30 degrees with respect to the T > C axis, matrix size = 64×72, Z-shim = −1.4).

### Preprocessing

Data preprocessing followed the procedure described in Morales et al. (2018): Imaging analysis was performed using SPM12 (Statistical Parametric Mapping; www.fil.ion.ucl.ac.uk/spm). The first five volumes of each run were discarded to allow for T1 stabilization. Functional images were realigned and unwarped using local field maps (Andersson et al., 2001) and then slice-time corrected (Sladky et al., 2011). Each participant’s structural image was segmented into gray matter, white matter, CSF, bone, soft tissue, and air/background images using a nonlinear deformation field to map it onto template tissue probability maps (Ashburner and Friston, 2005). This mapping was applied to both structural and functional images to create normalized images in Montreal Neurological Institute (MNI) space. Normalized images were spatially smoothed using a Gaussian kernel (6 mm FWHM). We set a within-run 4 mm affine motion cutoff criterion.

To extract trial-wise activation estimates, we used SPM to fit a design matrix to the preprocessed images. The design matrix included a regressor for each experimental trial, as well as nuisance regressors for instruction screens and physiological parameters. Trials were modeled as 33 millisecond boxcar functions, locked to the presentation of the stimulus, and convolved with a canonical hemodynamic response function. Trial-wise beta estimates were then used in multivariate analysis.

### Multivariate analysis

Only correct trials were used for decoding (75% and 76% of trials from included blocks in the detection and discrimination tasks, respectively). We chose to limit our decoding analysis to correct trials in order not to conflate effects of subjective confidence with those of objective accuracy, or stimulus type. However, we found that qualitatively similar results are obtained when analyzing all trials (unthresholded whole brain maps are available in this study’s NeuroVault collection: neurovault.org/collections/9912/).

Stimulus presence (present vs. absent) was decoded during detection blocks, and stimulus identity (clockwise vs. anticlockwise orientation) during discrimination blocks. Both decoding analyses used an LDA (Linear Discriminant Analysis) classifier with leave-one-run-out cross-validation and a searchlight radius of 4 voxels (∼257 voxels per searchlight). Significance testing was done using permutation testing to generate the empirical null-distribution. We followed the approach suggested by Stelzer, Chen, & Turner (2013) for searchlight MVPA measurements which uses a combination of permutation testing and bootstrapping to generate chance distributions for group studies. Per participant, 25 permutation maps were generated by permuting the class labels within each run. Group-level permutation distributions were subsequently generated by bootstrapping over these 25 maps, i.e. randomly selecting one out of 25 maps per participant. 10000 bootstrapping samples were used to generate the group null-distribution per voxel and per comparison. *P*-values were calculated per searchlight or ROI as the right-tailed area of the histogram of permutated accuracies from the mean over participants. We corrected for multiple comparisons in the searchlight analyses using whole-brain FDR-correction. A cluster-extent threshold was applied, ensuring that voxels were only identified as significant if they belonged to a cluster of at least 50 significant voxels (Dijkstra, Bosch, & van Gerven, 2017).

## Results

### Decoding of stimulus presence and orientation

We first searched for multivariate activation patterns that encoded information about stimulus orientation (in discrimination) and stimulus presence/visibility (in detection). In a whole-brain searchlight analysis, stimulus orientation could be reliably decoded only from the visual cortex (Fig. 2C). In contrast, information about stimulus presence was identified in parietal and prefrontal brain regions, including the dorsolateral prefrontal cortex, the middle frontal gyrus, and the precuneus (see Fig. 2A, for unthresholded classification maps, see neurovault.org/collections/9912).

**Figure 2:**
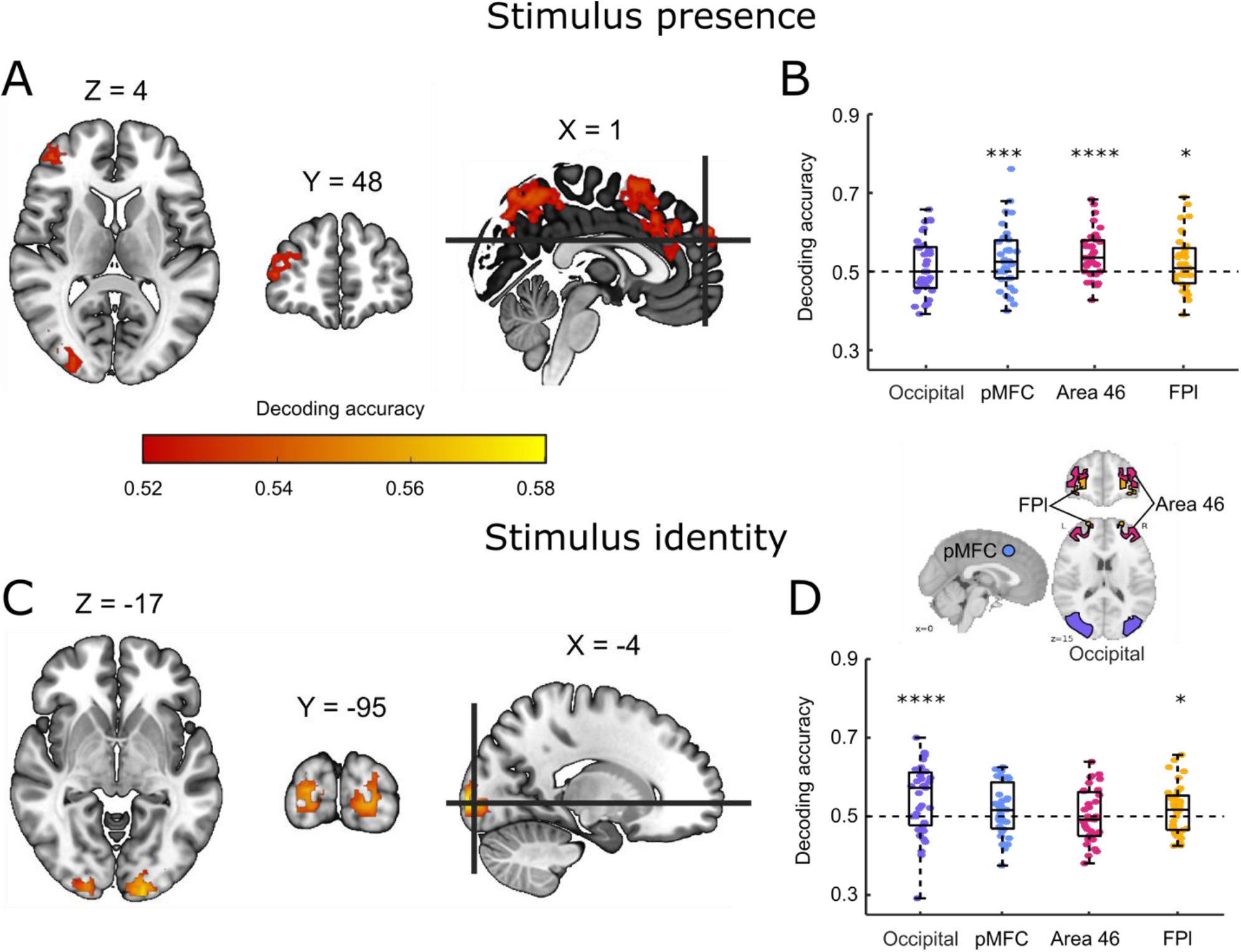
Decoding of stimulus presence and stimulus identity. A: whole brain searchlight decoding of stimulus presence versus absence in the detection task, correct responses only. B. decoding of stimulus presence versus absence in the occipital, pMFC, BA 46 and FPl ROIs. C: Whole brain searchlight decoding of stimulus identity in the discrimination task, correct responses only. D: decoding of stimulus identity in the four ROIs. Whole-brain maps are corrected for multiple comparisons at the voxel level with a cluster-size cutoff of 50 voxels. *: p<0.5, **: p<0.01, ***: p<0.001, ****<0.0001.

Based on these maps, we decided to focus our subsequent analyses on four regions of interest (ROIs): an occipital ROI, defined using the AICHA atlas as ‘occipital mid’ regions (Joliot et al., 2015) and three prefrontal ROIs which were also used in Mazor et al (2020): posterior medial frontal cortex (pMFC; an 8 mm sphere around MNI coordinates [0, 17, 46]), Brodmann area 46, and lateral frontopolar cortex (BA46 and FPl; both defined based on a connectivity-based parcellation; Neubert et al., 2014). Bilateral ROIs were defined as the union of the right and left hemispheres.

Within these four ROIs, stimulus orientation could be decoded significantly from occipital (*M* = 0.54, *SD* = 0.09, *p* < 0.0001) and FPl ROIs (*M* = 0.51, *SD* = 0.06, *p* = 0.04). In contrast, stimulus presence could be decoded from pMFC (*M* = 0.53, *SD* = 0.08, *p* = 0.0009), area 46 *(M* = 0.54, *SD* = 0.06, *p* < 0.0001) and FPl ROIs (*M* = 0.52, *SD* = 0.07, *p* = 0.015), but not from the occipital ROI (*M* = 0.51, *SD* = 0.07, *p* = 0.11). Classification accuracy showed a significant ROI x task interaction (*F*(3,32) = 5.31, *p* = 0.004; see Fig. 2, right panel), suggesting that stimulus presence (Fig. 2B) and stimulus identity (Fig. 2D) are encoded differentially across ROIs. Post-hoc contrasts revealed a significantly higher classification accuracy for detection compared to discrimination in area 46 (t(34)=3.06, p<0.005), with no significant difference between detection and discrimination decoding in the FPl, pMFC, or occipital ROIs.

### Behavioural analysis and confidence-based decoding

As previously reported in Mazor et al. (2020), task performance was similar for detection (75% accuracy, d’=1.48) and discrimination (76% accuracy, d’=1.50). Repeated measures t-tests failed to detect a difference between tasks both in mean accuracy (t(34) = −0.90, p=0.37, d=0.15, BF01 = 5.15), and d’ (t(34) = −0.30, p=0.76, d=0.05, BF01=7.29), indicating that performance was well matched. Within detection, participants were significantly more confident in ‘yes’ responses (mean confidence = 5.03 on a 1-6 scale) compared to ‘no’ responses (mean confidence = 4.21; t(34)=5.83, p<0.001, d=1.00). In contrast, confidence in discrimination ‘clockwise’ responses (mean confidence =4.28) was not significantly different from confidence in discrimination ‘anticlockwise’ responses (mean confidence =4.25; t(34)=0.31, p=0.76, d=0.05).

This absence of a significant difference between confidence in discrimination responses may indicate that a typical participant rated confidence similarly for discrimination ‘clockwise’ and ‘anticlockwise’ responses. Alternatively, it may be that some participants showed a bias towards higher confidence in ‘clockwise’ responses and others showed a bias towards higher confidence in ‘anticlockwise’ responses. Deciding between these two alternatives is important for interpreting our multi-voxel pattern analysis of discrimination responses: if single participants were consistently more confident in one of the two discrimination responses, above chance classification accuracy for discrimination may still be driven by differences in decision confidence, even if such differences average out at the group level (Gilron et al., 2017).

To decide between these two alternatives, we trained and tested an LDA classifier to predict participants’ decisions from their confidence ratings only. We used the same leave-one-run-out cross-validation procedure as in our MVPA analysis. This was done separately for the two tasks and for each participant. Confidence ratings successfully predicted detection responses, in line with a difference in mean confidence between detection ‘yes’ and ‘no’ responses (M=0.65, t(34)=9.70,p<0.001, d=1.64). Importantly, an LDA classifier also separated discrimination responses based on decision confidence (M=0.57, t=6.25, p<0.001, d=1.06), but to a lesser extent than in detection (t(34)=3.88, p<0.001, d=0.67 for a paired t-test testing the difference in classification accuracy between detection and discrimination). These analyses further emphasise the need to control for confidence when interpreting above-chance classification of detection and discrimination responses in higher-order brain regions in our data, as these may reflect person-specific differences in mean confidence between the two responses. Our next set of analyses was designed to control for this potential confound.

### Confidence-matching via downsampling

Prefrontal decoding of stimulus presence is consistent with the proposal that subjective visibility is represented in a frontoparietal network. However, it is also plausible that prefrontal decoding of detection reflects representations of confidence, instead of visibility. This alternative interpretation is in line with the finding that activity in prefrontal cortex is sensitive to variation in confidence (Vacarro & Fleming, 2018), and with our observation that confidence varied between detection decisions more than between discrimination decisions.

In our next analysis we therefore set out to determine whether our prefrontal ROIs would continue to represent stimulus presence *after controlling for decision confidence*. Having trial-wise confidence ratings allowed us to perfectly match not only mean confidence, but the entire distribution of confidence ratings for target present and target absent responses, and quantify the effect this had on classification accuracy. This was achieved by downsampling: for each participant and for each task, we selectively deleted trials until the two response categories had an equal number of trials for each confidence level (see Fig. 3A, left histogram). For example, if a participant had 15 trials in which they gave a confidence rating of 6, out of which only 3 were target absent trials, we randomly deleted 9 target-present trials in which the participant gave a confidence rating of 6, resulting in an equal number of confidence-6 trials for each response category. By then applying our presence/absence decoding analysis to these downsampled\ data, we were able to obtain a “downsampled” decoding accuracy, which reflected the ability of a classifier to determine stimulus presence vs. absence from activation patterns, after removing differences in confidence.

**Figure 3:**
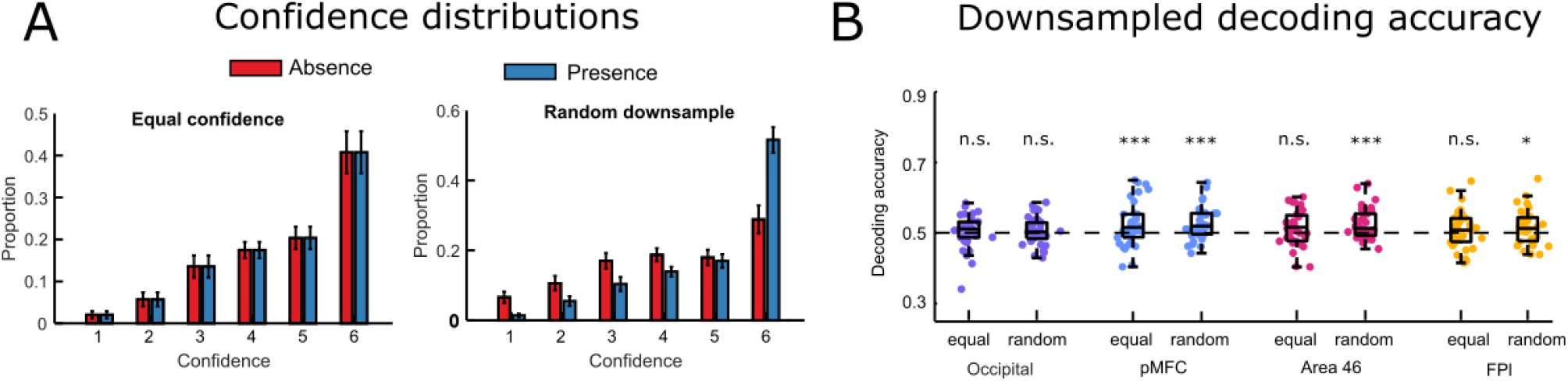
Stimulus presence downsampling analysis. A: for each participant, trials were deleted until confidence distributions were matched for target present and target absent responses. As a control analysis, we repeated this procedure with random downsampling, deleting the same number of trials irrespective of confidence ratings. B: presence/absence classification accuracy in the four ROIs for the equal confidence and random downsampling datasets. *: p<0.5, **: p<0.01, ***: p<0.001, ****: p<0.0001

To make sure any change in decoding accuracy was not simply due to a reduction in trial number, we also repeated this procedure with random instead of confidence-based downsampling, resulting in a second ‘random downsampled’ decoding accuracy value for each ROI. Importantly, this procedure of random downsampling ensures that the trial numbers in the two classes are the same as in the equalized confidence analysis, while keeping any confidence differences intact (see Fig. 3A, right histogram). Because there are multiple ways in which a dataset could be downsampled, for both types of analyses we repeated the procedure 25 times to take into account the variance created by selective sampling and then averaged decoding accuracy over these different downsampled sets. Finally, for statistical testing we created null distributions by following the same downsampling procedure on label-shuffled datasets.

When equalizing confidence, classification accuracy for decoding stimulus presence remained significant in pMFC (*M* = 0.52, *SD* = 0.06, *p*=0.002). However, decoding was no longer significant after equalizing the confidence distributions in FPl (*M* = 0.51, *SD* = 0.05, *p* = 0.11), and only marginally significant in area 46 (*M*=0.51, *SD*=0.05, *p*=0.07). In both regions, decoding was still significant after random downsampling (FPl: *M* = 0.52, *SD* = 0.05, *p*=0.02; area 46: M=0.53. SD=0.04, p=0.0017). A decrease in classification accuracy after equalizing confidence relative to random downsampling was marginally significant in area 46 (t(34)=-1.733, p=0.09, d=0.29), but not in the FPl ROI (t(34)=-1.615, p=0.11, d=0.27). In the pMFC ROI, classification accuracies for the confidence-matched and random downsamples were highly similar (0.524 and 0.525, t(34)=-0.20, p=0.84). Taken together, these results show that in pMFC, but not area 46 and FPl, stimulus presence/visibility can be reliably decoded independent of differences in decision confidence.

When decoding stimulus identity in the discrimination task, confidence-matching had no effect on classification accuracy relative to random downsampling (downsampled classification accuracy in the occipital ROI: M= 0.55, SD = 0.07; FPl: M = 0.52, SD = 0.06; pMFC: M = 0.51, SD = 0.06; area 46: M = 0.51, SD = 0.05, all pairwise comparisons with non-downsampled accuracy p > 0.28). This is consistent with there already being little difference in the (behavioural) confidence distributions between the two response types in discrimination blocks. Importantly, in pMFC, we observed no significant classification of stimulus identity, regardless of whether the analysis used confidence-matched data or not (downsampled classification accuracy: M=0.52, SD=0.06, p=0.2). In other words, in this prefrontal ROI, we were able to decode visibility (independently of confidence) but not identity.

## Discussion

What role the prefrontal cortex plays in visual awareness is much debated (e.g. Aru, Bachmann, Singer & Melloni, 2012; Boly et al., 2017). Here, we investigated whether prefrontal areas encode the visibility of a faint stimulus independently of stimulus identity and decision confidence. We first showed that a subset of prefrontal ROIs (pMFC and area 46) tracked stimulus presence during a detection task but not stimulus identity during a discrimination task, consistent with prefrontal involvement in encoding of stimulus visibility. Furthermore, classification accuracy was significantly higher for stimulus presence than for stimulus identity in area 46. However, because seeing a stimulus is associated with higher confidence than not seeing a stimulus, this asymmetry could also reflect confidence coding in frontal areas. To investigate this possibility, we tested whether decoding of stimulus presence remained significant after controlling for differences in confidence. We found that such decoding was indeed still possible in pMFC, but not in area 46. Taken together, these results suggest that pMFC, in contrast to area 46, encodes stimulus visibility over and above decision confidence. Furthermore, pMFC, unlike occipital regions, did not significantly encode stimulus identity, either when allowing confidence to freely vary, or when controlling for confidence in a downsampling analysis.

However, it is important to note that the interpretation of a “pure visibility” signal in pMFC is nuanced by a lack of significant difference between classification accuracies for stimulus presence and identity in this region. In other words, while we can decode stimulus visibility but not identity in pMFC, we cannot conclude that the decoding of these two quantities are themselves significantly different. Therefore, one viable alternative interpretation of our results might be that pMFC encodes a low-dimensional projection of rich perceptual input onto a decision axis: one that separates clockwise from anticlockwise gratings in discrimination blocks, and noise patches with and without a grating in detection blocks. Nevertheless, regardless of the nuance required when interpreting results in individual prefrontal ROIs, our results make clear that what may appear to be neural signatures of visibility in prefrontal cortex (e.g. in whole-brain searchlight decoding, such as in Figure 2) may on closer inspection be more closely related to differences in decision confidence.

Conceptually, visibility and decision confidence appear similar. They can both be defined in terms of precision: the precision of a visual percept in the first case, and the precision with which a decision is made in the second (Denison et al., 2017). Empirically, neural correlates of visibility and decision confidence overlap, specifically in the dorsolateral prefrontal cortex (dlPFC) but also in medial prefrontal, parietal, and insular cortices (Vacarro & Fleming, 2018).

Notwithstanding this conceptual and empirical overlap, visibility and confidence are not one and the same thing. Critically, within a Bayesian framework, decision confidence is defined as the probability correct of a particular response, and should therefore be sensitive not only to the precision of sensory representations, but also response requirements (Pouget et al., 2016; Bang & Fleming, 2018). Accordingly, visibility judgments scale with stimulus contrast even in trials in which participants make erroneous decisions, but confidence judgments show a different profile, and are sensitive to stimulus contrast only for correct responses (Rausch and Zehelteiner, 2016).

Despite a theoretical distinction between confidence and visibility, neuroimaging findings of visual awareness have often not been able to separate their respective contributions to differential brain activation. For example, it has not been possible to determine whether the dorsolateral prefrontal cortex is more active on aware versus unaware trials because it is sensitive to subjective visibility, or because participants are generally more confident in their decisions when they are aware of a stimulus. In an exploratory analysis of existing imaging data, we found that an apparent encoding of stimulus visibility in area 46 and lateral frontopolar cortex disappeared when controlling for subjective confidence. In contrast, pMFC encoding of visibility remained significant.

As reported in Mazor et al. (2020), univariate analysis of this data indicated a similar parametric modulation of confidence for detection and discrimination responses in pMFC. Specifically, a similar modulation of confidence in decisions about target presence and absence indicate that univariate signal in this region also scales with decision confidence. Univariate analysis did not reveal a pMFC modulation of visibility, which would manifest as an interaction of confidence and class in detection (because visibility is negatively correlated with confidence in ‘no’ responses, but positively correlated with confidence in ‘yes’ responses). However, a pre-registered cross-classification analysis revealed shared multivariate representations for discrimination confidence and detection responses indicating whether a stimulus is seen or not in pMFC and area 46 (Mazor et al., 2020; Appendix 8). We previously interpreted these findings as indicating that multivariate spatial activation patterns in area 46 and pMFC hold information about stimulus visibility, because like detection responses, confidence during discrimination might also track stimulus visibility (it is easier to determine what something is when you see it more clearly). Our current results corroborate this finding with respect to pMFC, and further show that above chance cross-classification in this region is not merely driven by differences in subjective confidence between ‘yes’ and ‘no’ responses during detection. Taken together, these results suggest that pMFC signal carries information not only about subjective confidence, but also about perceptual content, be it stimulus visibility, stimulus identity, or both.

Activation in pMFC is commonly found to correlate negatively with subjective confidence, or positively with uncertainty (Fleming, Huijgen & Dolan, 2012; Molenberghs et al., 2016; Vacarro & Fleming, 2018; Mazor, Friston & Fleming, 2020). In a recent study we found that univariate pMFC activation tracked the effect of decision difficulty, although it was insensitive to the precision of perceptual information in a motion perception task, which was instead tracked in posterior parietal regions (Bang & Fleming, 2018). Other work has shown that the pMFC is important for signaling when decisions or beliefs should be updated on the basis of new information (Fleming et al., 2018; O’Reilly et al., 2013). Novel paradigms may be necessary to further disentangle pMFC contributions to encoding stimulus visibility, and to relate this putative computational role to the encoding of other types of (perceptual and non-perceptual) uncertainty.

Our initial analysis specifically highlighted area 46 in the decoding of stimulus presence *without* controlling for confidence differences. This pattern of results is consistent with area 46 contributing to detection confidence, whereas more posterior prefrontal cortex (pMFC) may support visual detection responses, irrespective of differences in confidence. This result is in keeping with previous observations TMS to area 46 leads to lower overall perceptual confidence (a change in metacognitive bias), without affecting metacognitive sensitivity (Shekhar & Rahnev, 2018). In contrast, TMS applied to frontopolar cortex in Shekhar and Rahnev’s study led to increases in metacognitive sensitivity, without affecting confidence bias. We note that the contribution of prefrontal cortical subregions to visual metacognitive sensitivity (the coupling between confidence and accuracy) is difficult to assess using within-subject neuroimaging methods applied here as it requires modeling confidence noise across many trials. It remains to be determined whether the visual confidence signal in area 46 we observe here is specific to perceptual judgments (Lau & Passingham, 2006), or generalises to different task domains (Morales et al., 2018; Fleck et al., 2006).

Our results with respect to the lateral frontopolar cortex (FPl) are more difficult to interpret. We found that this area did not represent stimulus presence over and above confidence, but that it did represent stimulus identity, even after controlling for confidence differences between the different stimulus classes. Several factors may have contributed to these results. First, our observation that the FPl does not encode visibility irrespective of confidence does not mean that this region cannot play a role in visual awareness. In target absence trials, participants can sometimes be fully aware of the absence of a target – a case where visibility is low, but awareness (of absence) is high (Mazor & Fleming, 2020). Therefore, if FPl tracks content-invariant aspects of visual awareness, its activation may not differentiate between target presence and target absence. However, a representation of stimulus identity in FPl suggests that this area might also encode stimulus content. We are not aware of previous reports of decoding of visual content from the frontopolar cortex. Moreover, a recent meta-analysis reported no known effects of intracranial electrical stimulation of the frontopolar cortex on spontaneous reports of visual experience (Raccah, Block & Fox, 2021). Given the relatively modest effect sizes in FPl decoding of stimulus identity (M=0.51) in comparison to the more robust encoding of stimulus identity in occipital cortex (M=0.55), we are cautious in over-interpreting this surprising result. Future studies are necessary to explore to what extent FPl truly represents stimulus identity, and/or contributes to visual awareness.

Finally, when considering the implications of these findings for the study of visual awareness and its neural correlates, it is important to note the difference between subjective reports of stimulus awareness, and decisions about the presence or absence of a target stimulus in a perceptual detection task. While the first is a subjective decision about the contents of one’s perception, the second is a report of one’s beliefs about the state of the external world. Consequently, these two types of decisions draw on different sets of prior beliefs and expectations. For example, in detection, but not in subjective visibility reports, participants may adjust their decision criterion when noticing that they haven’t detected a stimulus in a long time. Furthermore, participants may base their detection responses not on the visibility of a stimulus, but on other visual and non-visual cues (adopting different *criterion content*; Kahneman, 1968). Our findings are based on the analysis of detection decisions, and their generalizability to reports of subjective awareness is an open empirical question.

To conclude, an exploratory data analysis revealed that stimulus presence could be decoded from prefrontal regions but that only the pMFC encoded stimulus presence after controlling for decision confidence. Future hypothesis-driven investigation is needed to replicate these exploratory results. We demonstrate the importance of controlling for confidence when investigating reports of awareness versus unawareness and propose a novel analysis approach to do so.

